# Spatially varying cell-specific gene regulation network inference

**DOI:** 10.1101/2025.10.17.683188

**Authors:** Yurui Li, Jin Chen, Haohan Wang

**Affiliations:** School of Information Science, University of Illinois at Urbana-Champaign, Champaign, IL 61820, USA; Department of Inflammation and Immunity, Lerner Research Institute, Cleveland Clinic, Cleveland, OH 44195, USA

**Keywords:** Gene Regulation Network, Deep Learning, Spatial Transcriptomics

## Abstract

Gene regulatory networks (GRNs), involving interactions between large numbers of genes, govern expression levels of mRNA and their resulting proteins to control cell functions. As many new sequencing technologies in single-cell and spatial resolutions are raised, the construction of GRNs gains opportunities to be generalized to the cell-specific level. Here we propose SVGRN, a deep learning model to infer cell-specific GRNs using spatial transcriptomics data. We model the gene expression, GRN matrix, and spatial coordinates of cells in the structural equation model (SEM) to learn gene interactions in an unsupervised way. Conditioned on the target cell position, the model is able to tune the whole tissue GRN to the target cell through also borrowing information from neighborhood cells. Results on simulated datasets show that SVGRN achieves better performance in cases with more noises, genes, or cells, which illustrates its ability to solve more complex situations. Our model was further applied to spatial transcriptomics datasets generated using different technologies and resolutions, including a seqFISH-based mouse embryo dataset and two Visium-based datasets from human cutaneous squamous cell carcinoma (cSCC) and fallopian tube tissues. The GRNs predicted by SVGRN from these datasets reveal dynamic gene regulatory patterns in mouse organogenesis, tumor development and the functional organization of the fallopian tube, highlighting the efficiency and broad applicability of our model.

## 1. Introduction

The collaboration between chromatin, transcription factors and genes generates complex regulatory circuits that can be modeled as gene regulatory networks (GRNs) [51]. Unraveling these interactions plays a critical role in understanding the underlying regulatory crosstalk that drives many cellular processes and diseases [36]. Traditional GRN inference methods rely on bulk transcriptomics data [49,40] or experimentally validated regulatory events. However, bulk profiling obscures regulatory programs specific to individual cell types or states, as it aggregates data from diverse cells within a tissue sample. This limitation has been overcome by single-cell multi-omics and spatial transcriptomics technologies, which enable GRN inference across distinct cell types, differentiation paths, and conditions. The advent of these technologies has led to the development of novel computational methods that infer GRNs at an unprecedented resolution.

The current computational methods employing diverse statistical or machine learning methods aim to reconstruct more comprehensive and precise gene regulatory networks. Many methods are developed to infer common GRNs of all cells or a specific subpopulation of cells through utilizing single-cell multi-omics datasets [72,29] or time series data [25,76]. There are only a few methods that can extend this inference to cell-specific GRNs [74,73]. In CeSpGRN [74], it models the gene expression data using Gaussian Copula Graphical Model (GCGM) [43,67] to conduct cell-specific inference from single-cell multi-omics and spatial data. LocaTE [73] utilizes an information theoretic approach that leverages single cell dynamical information together with the geometry of the cell state manifold to infer cell-specific GRNs. Although these methods enable people to understand GRNs on a finer-grained level, they often need additional measurements besides single cell gene expression such as scATAC-seq and RNA velocity, which adds more difficulties to the inference. These measurements may also introduce additional noise as they may come from different experiments.

Therefore, in this paper, we propose a deep learning method, SVGRN, solely based on spatially resolved gene expression data to infer varying cell-specific GRNs along the space without other measurements or prior knowledge. Relying on the decisive role of cell coordinates to its GRN and utilizing useful information from neighborhood cells, we are able to refine the GRN of the whole tissue to the specific target cell by borrowing information from its spatial position and neighboring cells. We implemented the modeling of gene expression and cells’ layout through conditional variational autoencoder (CVAE), and to the best of our knowledge, SVGRN is the first method based on a deep learning model to construct cell-specific GRNs using spatial transcriptomic data. The introducing of deep learning allows modeling spatially varying GRNs in a more comprehensive nonlinear framework and benefits from its computational power. Compared with CeSpGRN on several simulated datasets with various settings, SVGRN demonstrates advantages on more complex inference situations and maintains affordable running time on a larger number of genes. We applied our model to a cell-based spatial transcriptomics dataset of mouse embryos, demonstrating its ability to capture regional gene interactions during early organogenesis. Additionally, the predicted GRNs from the spot-based cSCC and fallopian tube dataset align with existing studies and reveal the functional tissue niches in space, further highlighting the broad applicability of SVGRN and its stable performance across different spatial transcriptomics technologies.

## 2. Methods

### 2.1 Gene interaction modeling with cell spatial coordinates

The underlying framework of SVGRN is derived from structural equation modeling (SEM), which is inspired from DeepSEM [62] using SEM for GRN modeling. SEM is a multivariate, hypothesis-driven technique that is based on a structural model representing a hypothesis about the causal relations among several variables [64]. In the context of GRN inference from spatial transcriptomics, these variables are the genes with available expression data on which we want to explore their interaction relationship, and the hypothetical causal relations are based on the regulation mechanism between genes. The structural model of SEM formulates each gene’s expression value as a function of the expression levels of other genes, enabling to capture the dependencies between genes in GRNs showing which genes influence others and to what extent.

In our case of GRN inference problem, let **X** ∈ ℝ^*n×m*^ be the gene expression matrix with *n* cells and *m* genes and **W** ∈ ℝ^*m×m*^ be the GRN adjacency matrix that represents the dependencies among different genes. The relationship between **X** and **W** can be formalized following the basic linear SEM [62] as

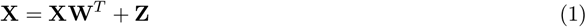

where **Z** ∈ ℝ^*n×m*^ is a matrix following Gaussian distribution representing the noise introduced during the transcription process. It helps to capture the unobserved variability or measurement errors in the spatial transcriptomics data, providing flexibility and robustness to the model. The equation (1) can be modified to

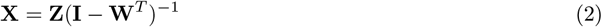

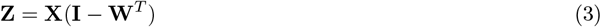

While linear SEM can provide basic gene dependencies, the actual gene regulation mechanisms in biological data are often more complicated. In transcriptomics data, the gene interactions usually involve diverse combinational and context-dependent factors such as external environmental response and cell-cell signaling, which makes the linear model challenging to capture this complexity. Therefore, we transform equation (2) and (3) to a nonlinear version of the SEM following DAG-GNN [71] to fully utilize the computational ability of deep learning

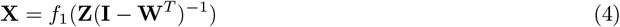

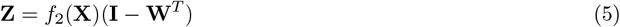

where *f*_1_ and *f*_2_ are parameterized functions performing nonlinear transforms on **Z**(**I** *−* **W**^*T*^)^*−*1^ and **X**, respectively.

When considering cells in spatially organized environments, their behavior is significantly influenced by spatial context factors. Depending on where a cell is located, specific genes will be turned on or off due to signals received from neighboring cells, effectively dictating its development and function based on its spatial context [57]. In the developing embryo, positions of cells within the concentration gradients of small diffusible molecules called morphogens can also trigger different gene regulation patterns [33]. Spatial transcriptomics, capturing transcriptional activity at distinct spatial locations, provides opportunities for us to include the positional context effects in our model. By incorporating spatial coordinates, the model can take into account how a cell’s location affects its gene expression, allowing it to capture microenvironmental effects that shape cellular function and differentiation. Accordingly, based on equation (4) and (5), we further bring **Y** ∈ ℝ^*n×*2^, the spatial coordinates of cells, into the model. The updated equations will become

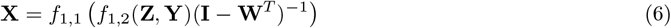

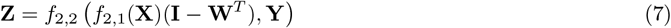

where *f*_1,1_, *f*_1,2_, *f*_2,1_, *f*_2,2_ are also nonlinear parameterized functions and implemented as multilayer neural networks in Section 2.2. Including **Y** in the functions *f*_1,2_ and *f*_2,2_ enables the model to learn the nonlinear dependencies between gene expression and the spatial organization of cells, while simultaneously capturing gene regulatory relationships. This design preserves the general symmetry between these two equations, providing a structured and balanced representation of spatial effects. Furthermore, this symmetry aligns with the architecture of the conditional variational autoencoder, making it well-suited for implementation within this framework, as we will further elaborate in Section 2.2.

Involving coordinates in the SEM establishes a bridge between gene expression and spatial location information. We can not only capture the gene-gene dependency from the gene expression data, but also consider the contribution to GRN from the spatial layout of cells. This provides the opportunity to extend the GRN on the whole tissue to single cells.

### 2.2 SVGRN framework based on CVAE

To integrate both equations (6) and (7), we use a conditional variational autoencoder (CVAE) [63] to model the relationship among gene expression matrix, GRN matrix, and cell coordinates. The CVAE structure consists of two components, a encoder compressing the input data into a latent representation and the decoder reconstructing the original data from this latent space, conditioned on additional input information. The encoding and decoding processes can be respectively aligned to the modeling in equations (7) and (6), and they are implemented using neural network structures to apprehend the non-linear relationships in data. Given the gene expression ***X*** and coordinates ***Y*** of cells from spatial transcriptomics, the encoder is trained to learn the distribution *q*_*ϕ*_(***Z***| ***X, Y***), parameterized by *ϕ*, which maps them into the latent space as ***Z***. This enables capturing how spatial information and gene expression are interrelated, resulting in a compressed representation of this complexity. Functioning in the opposite way, the decoder *p*_*θ*_(***X***| ***Z, Y***), parameterized by *θ*, reconstructs the gene expression ***X*** based on the latent representation ***Z*** and the spatial coordinates ***Y***. It models how gene expression varies as a function of both the regulatory signals encoded in the latent space and the spatial locations of cells, effectively implementing the SEM model in a non-linear manner. For the GRN matrix **W** in the equations, it will serve as part of the parameters in both encoder and decoder, and be learned through training.

The more detailed neural network structure of SVGRN is shown in Fig. 1, which also follows the functions in our SEM equations. 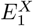 corresponding to *f*_2,1_(*·*) in equation (7) is the neural network to encode gene expression, and we additional add 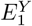 for cell coordinate encoding to better capture their spatial patterns. The gene expression encoder 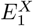 is followed by a GRN layer to directly capture gene-gene dependency from gene expression embeddings. *E*_2_, implementing *f*_2,2_(*·*), combines expression embeddings with coordinate features to further project them to latent space. With a symmetrical structure, *f*_1,1_(*·*) and *f*_1,2_(*·*) are respectively implemented as neural network decoders *D*_1_ and *D*_2_. *D*_2_ decodes the features of latent variables and coordinate features, followed by a reverse GRN layer to remove the gene dependency. Then *D*_1_ is used to finally recover the input gene expression. To let GRN layers fully comprehend interactions between genes, we avoid involving other combinations across genes in the rest of neural network structures. Therefore, each encoder or decoder shares the same weight across all the genes and only conducts calculations on each single gene.

**Fig. 1.**
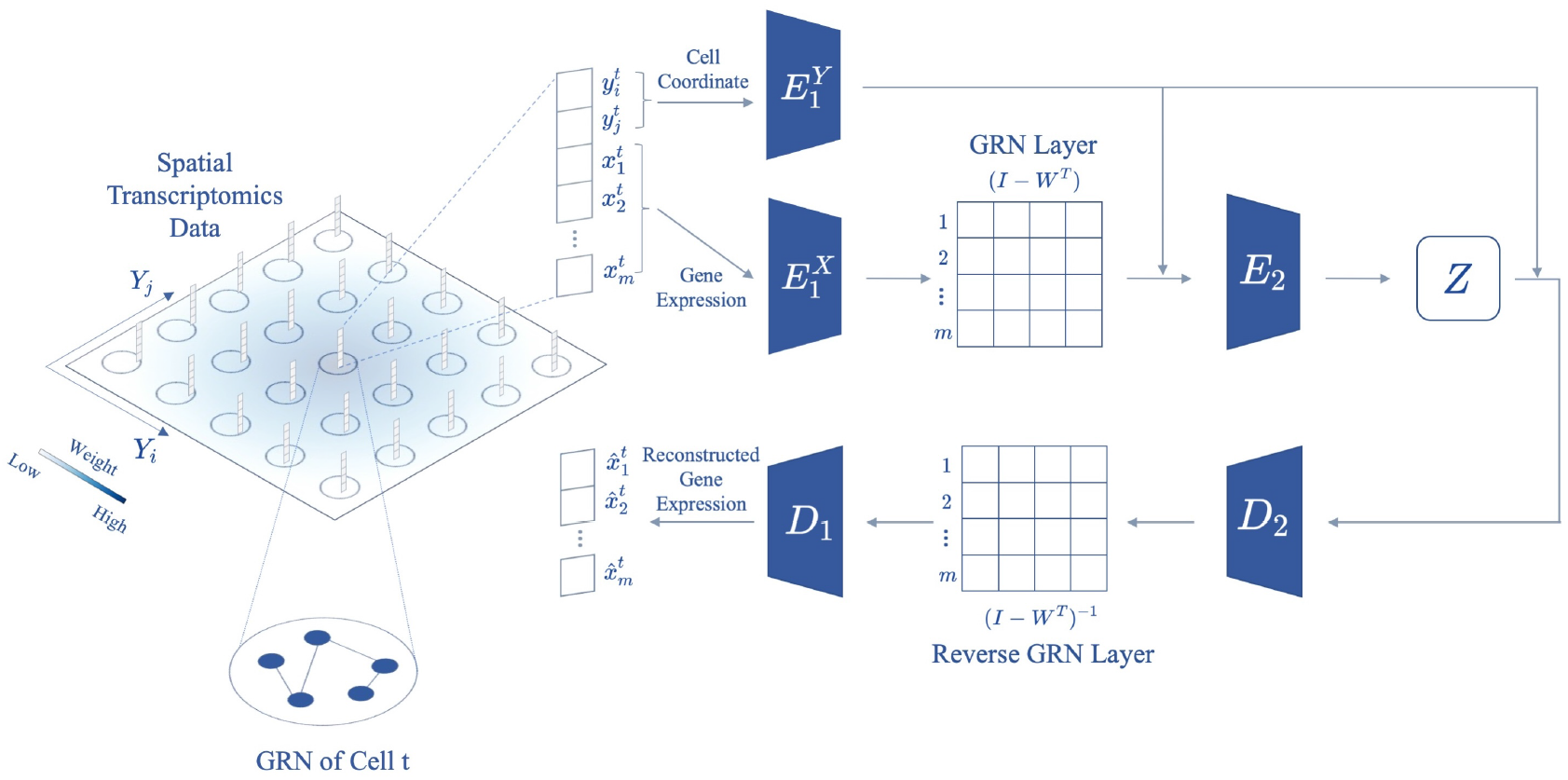
The SVGRN model architecture. The model takes the gene expression and its corresponding spatial coordinate for each spot as input from spatial transcriptomics data. 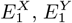, and *E*_2_ encode input data into latent space, and *D*_1_ and *D*_2_ are decoders to reconstruct the gene expression data. The GRN layer and inverse GRN layer learn through training to capture gene interactions.

Given the dataset 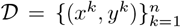 where *x*^*k*^ = **X**_*k*,:_ and *y*^*k*^ = **Y**_*k*,:_, the goal of CVAE is to fit a model of the conditional probability distribution *p*(*x* | *y*). It learns to recover the gene expression of each cell conditioned on its spatial coordinate and therefore, relates cell-specific gene expression patterns with their specific places in the cell layout. The loss function of SVGRN extends the loss function for CVAE with an additional *L*_1_ norm to regularize the adjacent matrix **W**, which is defined as follows

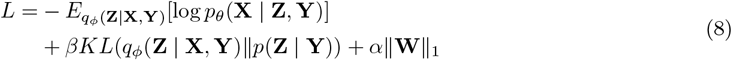

where E is the expected value function, KL denotes to KL-divergence function, and *α* and *β* are hyperparameters. Applying *L*_1_ regularization encourages sparsity in the learned GRN matrix **W**, reflecting the reality that only a subset of genes directly regulate each other in gene regulatory networks.

### 2.3 SVGRN training for cell-specific GRNs

The training of our SVGRN model can be devided into two stages, which gradually refine the learning from all cells in the tissue to a specific target cell. The pseudocode of this two-stage training is illustrated in Algorithm 1. Stage 1 training establishes a general GRN by using gene expression and spatial coordinates of all cells, providing a foundational network that captures the tissue’s overall regulatory patterns. Stage 2 then refines this network by conditioning on the target cell’s position and leveraging information from neighboring cells to learn a GRN specific to the target cell’s microenvironment. This two-stage approach establishes a biologically meaningful hierarchy by first capturing tissue-wide regulatory signals and then focusing on cell-specific nuances driven by local interactions, enabling precise inference of spatially resolved GRNs at the single-cell level. The following paragraphs will provide further details on each training stage.

In many cases, we only have access to gene expression data and spatial information, with limited prior knowledge of TF-gene interactions. Obtaining such prior knowledge often requires significant additional effort and may not always be applicable, as TF-gene interactions can vary depending on the tissue type, developmental stage, section position, and individual regulatory differences. To address these challenges, the first training stage is designed to avoid reliance on prior interaction data and instead learn a general GRN specific to the tissue being studied. Additionally, this stage incorporates spatial information to capture the relationship between gene expression and cellular coordinates. This allows the trained model to serve as a foundation for the second stage, where it can be fine-tuned to target specific cells by conditioning on the target coordinate.

In the first training stage, the training dataset 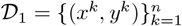 is composed of pairs of gene expression and corresponding coordinates for each cell *k*. The loss function follows the same form of equation (8) and can be further written as follows

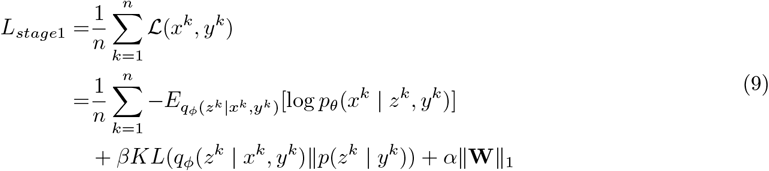

By training on all cells equally at this stage, the model learns a general distribution of gene expression of all cells given their spatial positions. In its GRN layers, the model starts with a randomly initialized GRN matrix and obtains an overall GRN representation on the entire tissue. It can serve as a good start for the following learning for each single cell.

In the second training stage, we further specialize our model to the single-cell level. Since in the previous stage, the model has already learned stable encoders and decoders, we only update the parameters of GRN layers and freeze all other layers in this stage. Here, we adjust training strategies based on two assumptions about the relationship between a cell’s spatial information and its gene expression profile. Firstly, we assume that the position of a cell can serve as a unique identifier or label for its GRN after the training in the first stage where the model learns the relationship between gene expression and its specific position across the dataset. By conditioning the model on these spatial positions, it captures the spatially-dependent variations in GRNs, allowing the model to further infer cell-specific GRNs of the target cells effectively based on the cell’s location. According to this assumption, when focusing on learning the GRN for a target cell *t*, we fix all the positions in the dataset as the position of this target cell. The training dataset of Stage 2 for target cell *t* will become 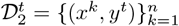. By doing so, the model is conditioned entirely on the target cell’s spatial context, enabling it to learn GRNs specific to the unique microenvironment of that cell. As we are training to model the distribution *p*(*x*| *y*^*t*^), the parameterized GRN matrix will be updated towards the GRN of cell *t* in the fixed position *y*^*t*^ which the whole model is conditioned on. This approach allows the model to focus on regulatory relationships specific to the target cell’s location, refining the general GRN from Stage 1 into a cell-specific network.

Secondly, we assume that the GRNs in the closer cells will tend to be similar, while cells that have a larger distance will have more differences in their GRNs, which is especially apparent in developing tissues. Through this assumption, when training the model to get the GRN for a target cell, we can not only use the single data for this target cell, but also be able to borrow information from its neighboring cells, as they could in some content provide some reference to the cell we are studying. In developing tissues such as embryos, GRNs exhibit smooth transitions across space due to the coordinated tissue development and gradual patterning along body axes. During embryogenesis, cell differentiation exhibit spatial continuity as they progressively transition from progenitors to specialized cell types [7,5]. This progression is often governed by spatially distributed gradients of morphogens, which gradually influence gene regulation patterns and expression throughout the tissue [6]. Consequently, cells that are spatially close, even if they belong to different types, often share similar GRNs.

When applied to fully developed tissues, although cells in mature tissues are more specialized, there are still some information that we can utilize from neighborhood cells to infer the target cell’s GRN as spatially close cells remain influenced by their shared microenvironment. Factors such as paracrine signaling, cell-cell interactions, and extracellular matrix-mediated communication contribute to maintaining localized regulatory similarities. While GRNs in mature tissues are generally less continuous than in developing tissues, spatial influences can still impose a degree of similarity in regulatory networks for cells in close proximity. This contextual information can be utilized to enhance the inference of a target cell’s GRN in mature tissues.

To implement this assumption, we introduce a kernel weight **K**_*kt*_ to the loss function, which shows how much information we can utilize from a cell *k* when focusing on a target cell *t* according to their similarity in spatial location. The kernel weight **K**_*kt*_ is calculated following the radial basis function kernel:

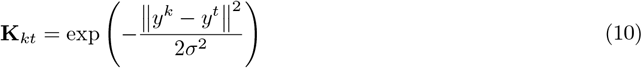

where *σ* is the hyper-parameter accounting for the range of neighboring cells that the target cell will refer to. The closer cells to the target cell *t* will have larger weights in loss values, while the weights of further cells will be much smaller. In this way, the neighborhood cells can have more contributions to model training for this target cell. By gathering the two assumptions the training stage 2 is based on, the loss function for this stage is defined as

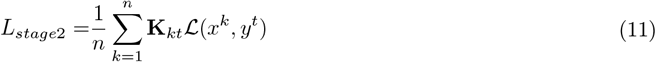

#### Algorithm 1

SVGRN Two-Stage Training for Cell-Specifc GRNs

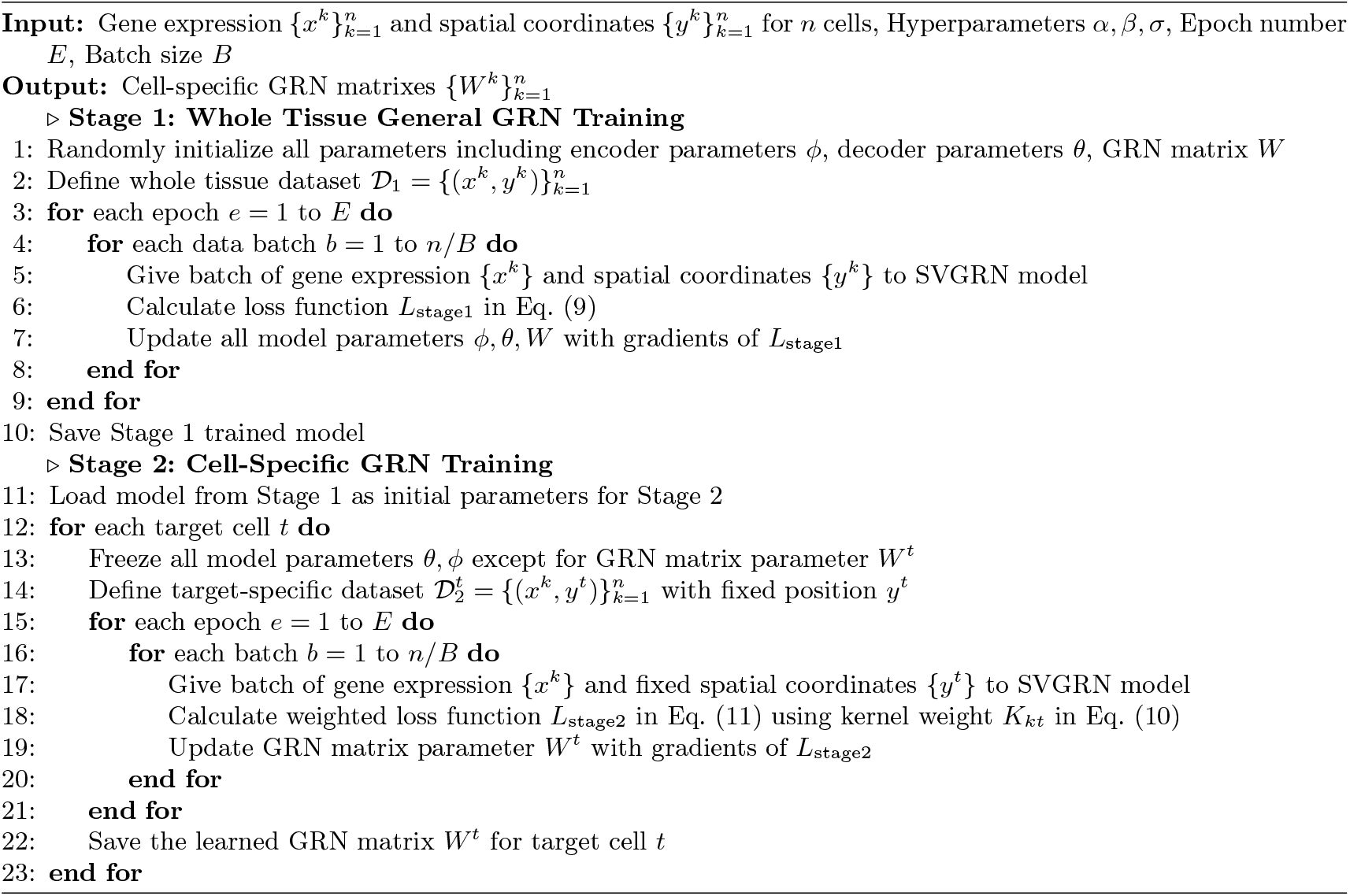

## 3 Results & Discussion

### 3.1 Experiments on simulation datasets

Due to the lack of datasets with known cell-specific GRNs, we evaluated our model’s GRN inference capabilities using simulated datasets generated by scMultiSim [41]. scMultiSim enables the simulation of scRNA-seq data given dynamic GRN across cells and the spatial organization of cells. To approximate real-world heterogeneous spatial transcriptomics data, we conducted a series of experiments, varying critical parameters such as gene numbers, cell numbers, and intrinsic noise levels, which are all crucial factors impacting GRN inference accuracy. In scMultiSim, the parameter *intrinsic*.*noise* adjusts the noise in the transcription process, where higher noise levels increase the difficulty of accurately inferring GRNs [41].

To benchmark our model’s performance in GRN inference, we adopted two metrics from the BEELINE framework: the early precision ratio (EPR) and the area under the precision–recall curve ratio (AUPRC ratio) [59]. These metrics provide insight into the accuracy of GRN constructions and indicate how well a model performs in predicting edges between genes. The early precision value measures the fraction of true positives among the top *k* predicted edges, with *k* set to the number of edges in the actual network. EPR represents the ratio of early precision value between the model and the random predictions, and AUPRC ratio is the ratio of the AUPRC between those.

#### SVGRN captures dynamic GRNs through cell-specific training

In Fig. 2, we show the visualization of cell layout in simulated dataset with 2000 cells, 110 genes and 0.1 intrinsic noise. Given an initial GRN matrix, scMultiSim can randomly pick pairs of genes to add new interactions, enlarge interaction strength, remove existing interactions, or weaken interaction strength to generate cell-specific GRNs cell by cell. To create a spatial layout where cells with similar GRNs are positioned close to each other, we used the “layers” parameter in scMultiSim for dataset simulation. This parameter controls placing cells one by one on a spatial grid, with new added cells more likely to be placed near existing cells with similar GRNs, forming a layered structure of GRN clusters. In Fig. 2 (a), we clustered cells by their GRNs and it demonstrates that the cell-specific GRNs follow a layered changing pattern across the space, which meets with our assumptions about closer cells having more similar GRNs.

**Fig. 2.**
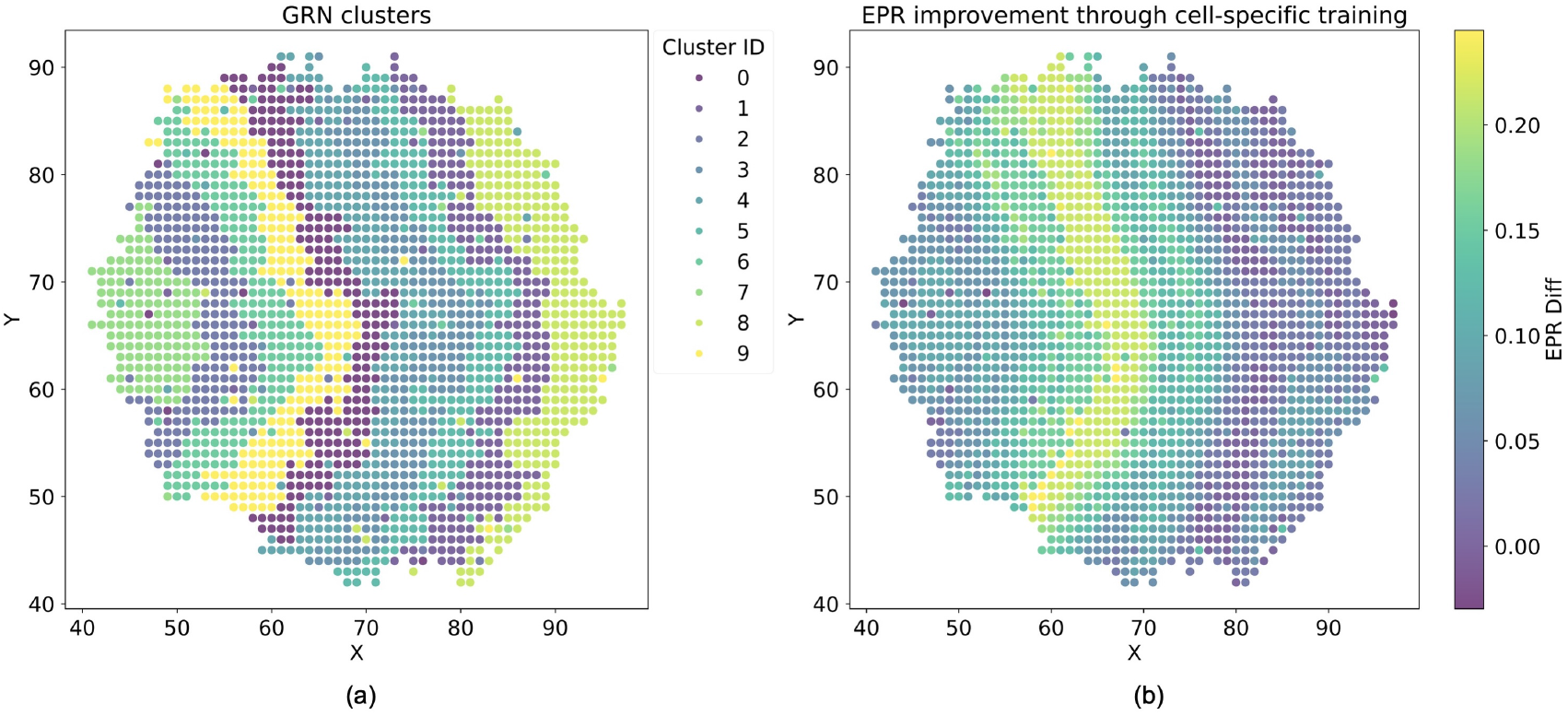
(a) The GRNs in all cells are clustered into 10 classes and the cluster ID for each cell is shown in their spatial layout. (b) For each cell, the difference between the initial EPR and that after the cell-specific training is calculated and shown as a heatmap.

With generated single-cell GRNs by scMultiSim, we calculated the EPR of each cell before and after the second training stage for cell-specific GRN refinement. The spatial distribution of EPR improvement values is shown in Fig. 2 (b). Results demonstrate that starting from a general whole-tissue GRN, our model effectively fine-tunes GRNs for each individual cell. Most cells show EPR improvements, validating the effectiveness of the cell-specific training stage. The pattern of EPR changes also aligns with the layered GRN distribution. The improvement values are similar in each layer shape and follow the real GRN cluster distributed pattern from left to right in the cell layout. It means that the model trained in the second stage can capture this layered distribution of cell-specific GRNs since GRNs in each layered cluster are similar and so that will have similar improvement when model is trained started from the same whole-tissue GRN to learn these cell-specific GRNs in all cells. This reflects the model’s ability to leverage spatial layout and nearby information, which enables the separate training on each cell to capture GRN variation across cells, leading to dynamic results in the space.

#### SVGRN shows advantages in more complex settings

We compared our model’s performance to CeSpGRN, a statistical method for inferring cell-specific GRNs from scRNA-seq data. Although LocaTE is also capable of inferring single-cell GRNs, we did not compare our model with it because it requires dynamic single-cell profiles, which is not well-suited to the context of our study. The average metrics across all cells are shown in Fig. 3, demonstrating that our model generally outperforms CeSpGRN, especially in cases with higher noise levels or larger numbers of genes and cells.

**Fig. 3.**
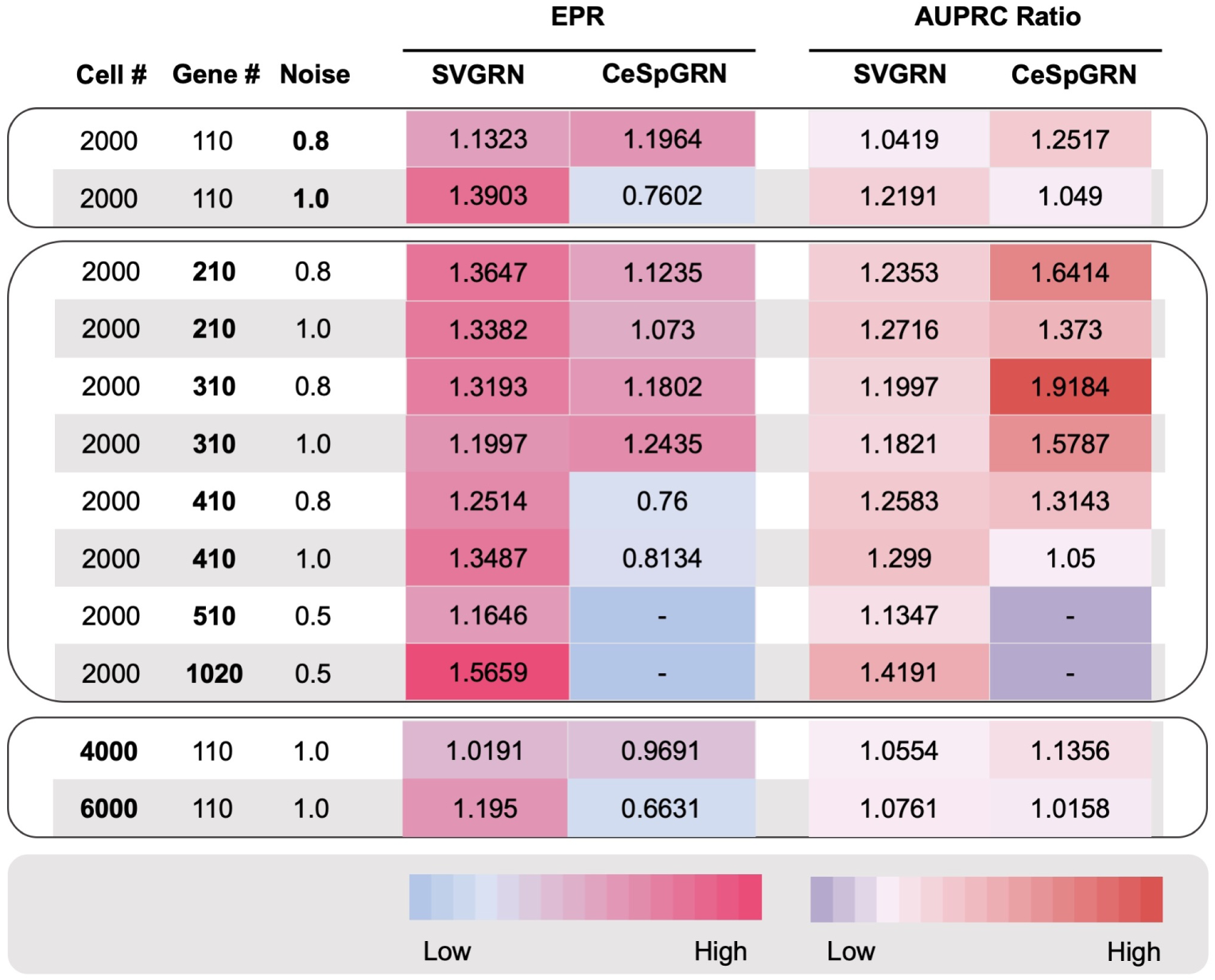
Result comparison of EPR and AUPRC ratio between SVGRN and CeSpGRN on simulation datasets with various cell numbers, gene numbers and noise levels. “-” represents unavailable results within affordable time spending.

Real biological datasets from single-cell sequencing often contain substantial noise due to technical factors such as low RNA capture rates [34], limited sequencing depth [4], and batch effects [58], along with biological variability across cell cycle phases [35]. Spatial transcriptomics introduces additional noise from extra steps that preserve spatial information during sequencing [68], such as unpredictable spatial noise from cell damage during cryosectioning and exposure to reagents for staining and mRNA release [15]. This noise can obscure subtle, meaningful interactions or amplify irrelevant signals, complicating accurate GRN construction. Thus, it is essential for our model to remain precise and reliable in noisy conditions. To assess this capability, we tested our model on simulated datasets with noise levels of 0.8, and 1.0. At a noise level of 0.8, our model achieved similar results to CeSpGRN, while at a noise level of 1.0, it demonstrated a clear advantage in both EPR and AUPRC ratios. These findings highlight SVGRN’s ability to infer underlying GRN patterns even in highly noisy data.

Moreover, in practice, tissues are composed of a vast number of diverse cells, contributing to complex tissue organization. A model that performs well with larger numbers of cells can better capture these complexities to deepen our understanding of cellular behavior. To evaluate this, we tested our model on datasets with increasing cell counts. For datasets with 4000 cells, our model showed higher EPR. It maintained stable performance as the cell number grew and got both higher EPR and AUPRC ratio in 6000 cells.

We also examined performance on datasets with increasing gene counts, which introduce more potential edges and a more complex interaction environment. As shown in Fig. 4, SVGRN, leveraging a deep learning framework, demonstrates superior computational efficiency. Using four GPUs for training, SVGRN’s runtime remains stable, while CeSpGRN’s runtime for datasets with 510 genes exceeded 48 hours, growing rapidly with he number of genes. We compared EPR and AUPRC ratios for both methods on datasets with 210, 310, 410, 510, and 1020 genes (Fig. 3). As it is hard to get the results of CeSpGRN on 510 and 1020 genes within affordable time spending, these results are marked as “-” in the figure. On datasets with 210, 310, 410 genes, our model achieved obvious higher EPR than CeSpGRN and still maintained stable results even with 1020 genes. This demonstrates SVGRN’s robustness in inferring GRNs when facing increasing cellular diversity and complex gene interactions.

**Fig. 4.**
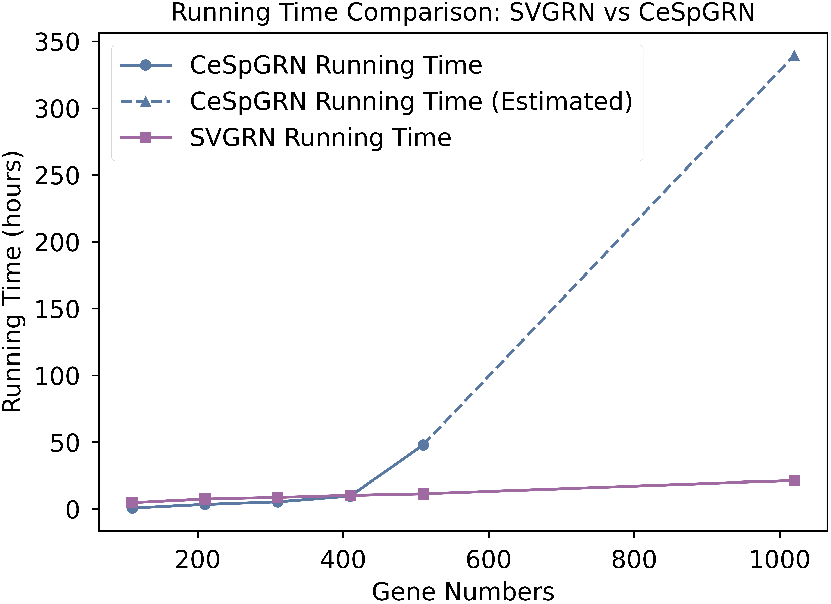
Running time comparison between SVGRN and CeSpGRN

### 3.2 Experiments on the cell-based spatial transcriptomics dataset of mouse embryos

Unlike simulated data, the biological environment is inherently complex, making it challenging to obtain true cell-specific GRNs through experiments. Using spatial transcriptomics data, the predicted cell-specific GRNs from our model provide a unique view of cell development within spatial contexts. Although some spatial transcriptomics technologies such as Visium [50] can preserve spatial locations of transcripts using NGS barcoding techniques, they typically measure transcriptomes in barcoded spots of 55μm in diameter, each containing about 1-10 cells, rather than achieving single-cell spatial resolution [52]. Some multiplexed single-molecule FISH (smFISH) technologies, such as seqFISH [46,45] and multiplexed error-robust FISH (MERFISH) [12], and in situ sequencing (ISS) methods [32] offer single-cell and single-molecule resolution. Compared to spot-based approaches with equidistant spatial data points, cell-based methods can better reflect true cellular distributions within tissues. This provides an opportunity to infer cell-specific GRNs in individual cells and incorporate more precise influences from neighboring cells based on actual spatial distances. In this section, we applied our model to a seqFISH-based spatial transcriptomics dataset [44] of mouse embryos at the 8–12 somite stage to infer GRNs. Early-stage embryonic tissues, such as those in this dataset, exhibit progressive sharpening of gene expression profiles as cells differentiate into specific identities [47]. Intermediate cellular states result in smooth transitions of gene expression and GRNs across spatial regions and cell types, aligning with our model’s assumption that closer cells share similar GRNs under the weight kernel in the second training stage.

We trained our model on sagittal section slice 2 of embryo 2, containing 6,880 cells and 351 genes from seqFISH. From the inferred cell-specific GRNs, we calculated the variance of each gene pair’s absolute edge weight across all cells to identify the most variable edges. Those edges represent gene interactions that fluctuate most along the space, potentially uncovering underlying regional gene regulation dynamics. From these edges, we can identify genes with variable functions in different embryo regions. Fig. 5 (a) demonstrates the gene appearance frequency in the top 100 most variable interacting gene pairs across cells. The most frequently shown genes include Hoxb9, Hoxd4, En1, Nepn etc., which all proved to have regional functions in mouse embryos [23,48,55,54,3,1].

**Fig. 5.**
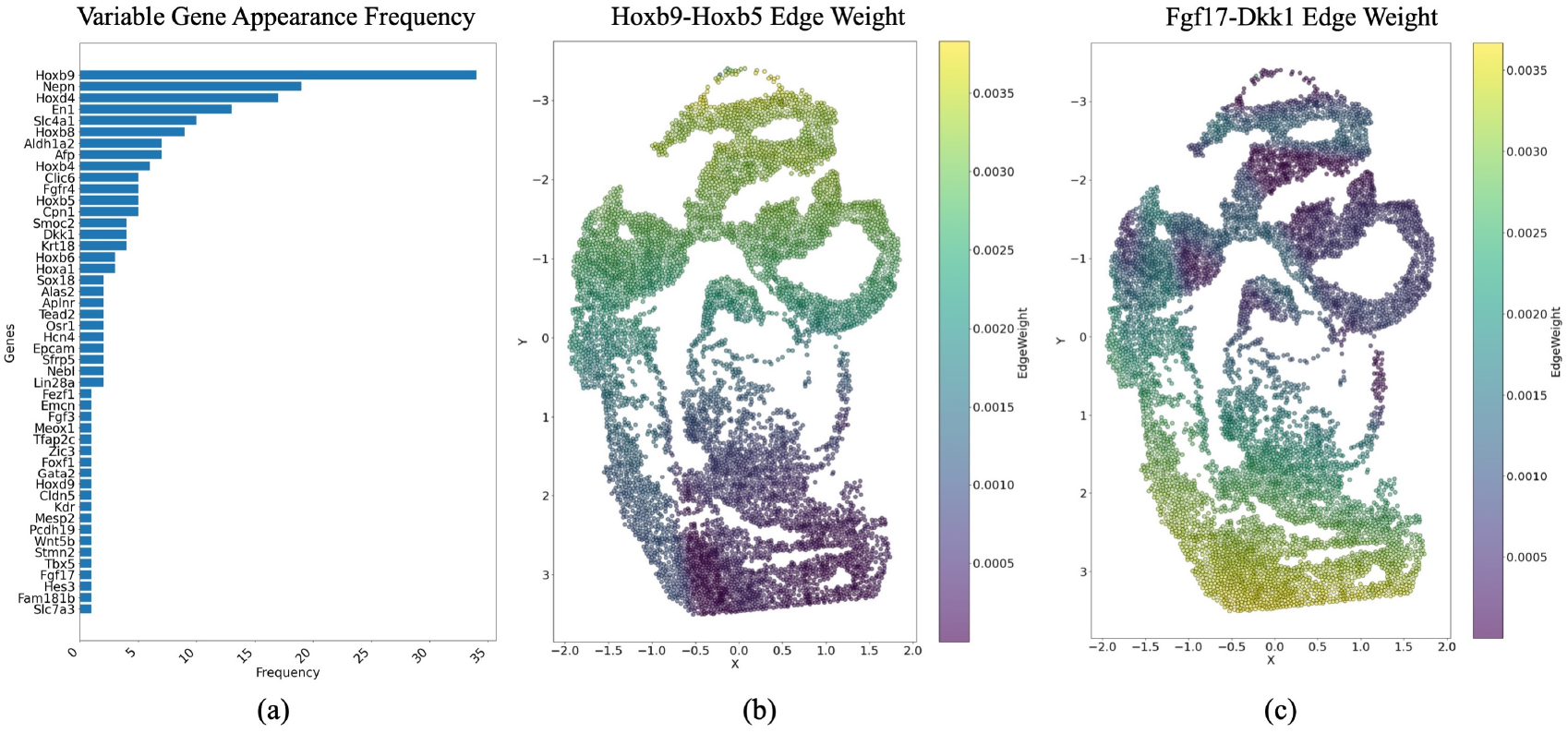
(a) The gene appearance frequency in the top 100 edges with highest edge weight variance value accross all the cells. (b) Hoxb9-Hoxb5 edge weight changing pattern in the mouse embryo tissue.(c) Fgf17-Dkk1 edge weight changing pattern in the mouse embryo tissue. In (b)-(c), largest values of X correspond to the tail region, and lowest values of Y correspond to the head region.

By analyzing all the inferred single-cell GRNs, we found that the model is able to capture some regional gene interactions functioning for different body segments of early-stage mouse embryos. Hox genes such as Hoxb9, Hoxd4, are key regulators of positional identity along the head-to-tail axis, determining body segment identities by controlling the expression of region-specific genes, thus shaping the vertebrae and limb morphology [23,48,55]. Hox genes can directly regulate the expression of other Hox genes within the same cluster or even across different clusters, functioning in overlapping and sequential domains [18,60]. This kind of interaction is particularly important for neural tube development, affecting the development of different neuronal populations and axial skeleton formation [2,14]. The gene interaction weight-changing pattern shown in Fig. 5 (b) illustrates possible interaction between Hoxb genes and their relationship with the neural tube development at the 8–12 somite stage of the embryo. The higher interaction appearing in the head region of the embryo shows the neural tube formation in the head region, where it then starts differentiating into brain regions and progresses toward the tail at this stage of development. Another pair of gene interactions that has a relationship with a specific region in mouse embryos is between Dkk1 and Fgf8 subfamily. Dkk1 may directly modulate the level of Wnt activity, thereby indirectly regulating Fgf expression [8] or this may suggest that Dkk1 acts over the Fgf pathway in a Wnt-independent fashion. More importantly, these interactions modulate cell movement inside the forming limb bud and Fgf8 activation in response to Dkk1 can be responsible for the observed limb phenotype [53]. Fig. 5 (c) demonstrates the captured interaction changing between Dkk1 and Fgf17 which belongs to the Fgf8 subfamily. It shows a relatively stronger interaction in the lumbar region which may show the merging of hindlimb buds located posterior to the developing forelimbs and close to the tail.

### 3.3 Experiments on the spot-based spatial transcriptomics dataset of cSCC

Compared to cell-based spatial transcriptomics data, spot-based data leads to a more complex case as each spot can contain multiple cells. Here, we further applied our model to a spot-based ST dataset of human cutaneous squamous cell carcinoma (cSCC) [27], which provides samples with both tumor and adjacent normal tissue regions. From the results, we demonstrate our model’s capability for spot-specific GRN inference in relatively mature tissue. Although each spot aggregates the expression of multiple cells, these cells are often spatially constrained and belong to related cell types or states. The aggregated transcriptomics data, therefore, reflects the dominant regulatory features of the region, which are also influenced by neighboring spots. It provides opportunities for our model to be used in this case that our model can still learn some useful information from the neighborhood spots and the coordinates of target spots. Our model helps to reveal how regulatory mechanisms gradually evolve from the tumor core to the leading edge and into adjacent normal regions in cancer. This capability is particularly valuable given the common lack of matched normal tissue from the same patient to be compared with tumors, which limits biological comparative experiments at high resolution. By generating GRNs tailored to individual spots in their spatial environment, our model supports the study of tumor development, tumor-local tissue crosstalk, and regulatory shifts that may drive tumorigenesis, offering insights into the spatially dynamic interactions underlying cell behavior and cancer progression.

In this dataset, we focused on the skin section from patient 2, which includes 17,139 genes across 1,933 spatial spots. The H&E stained tissue section of this patient is shown in Fig. 6 (d) as a reference of the locations of the tumor and normal regions which comes from this cSCC dataset [27]. For model training, we selected 666 spots located on the tissue, covering both tumor and normal regions, and used the top 40 marker genes of basal, cycling, differentiating, and tumor-specific keratinocyte (TSK) clusters shared between normal skin and cSCC.

**Fig. 6.**
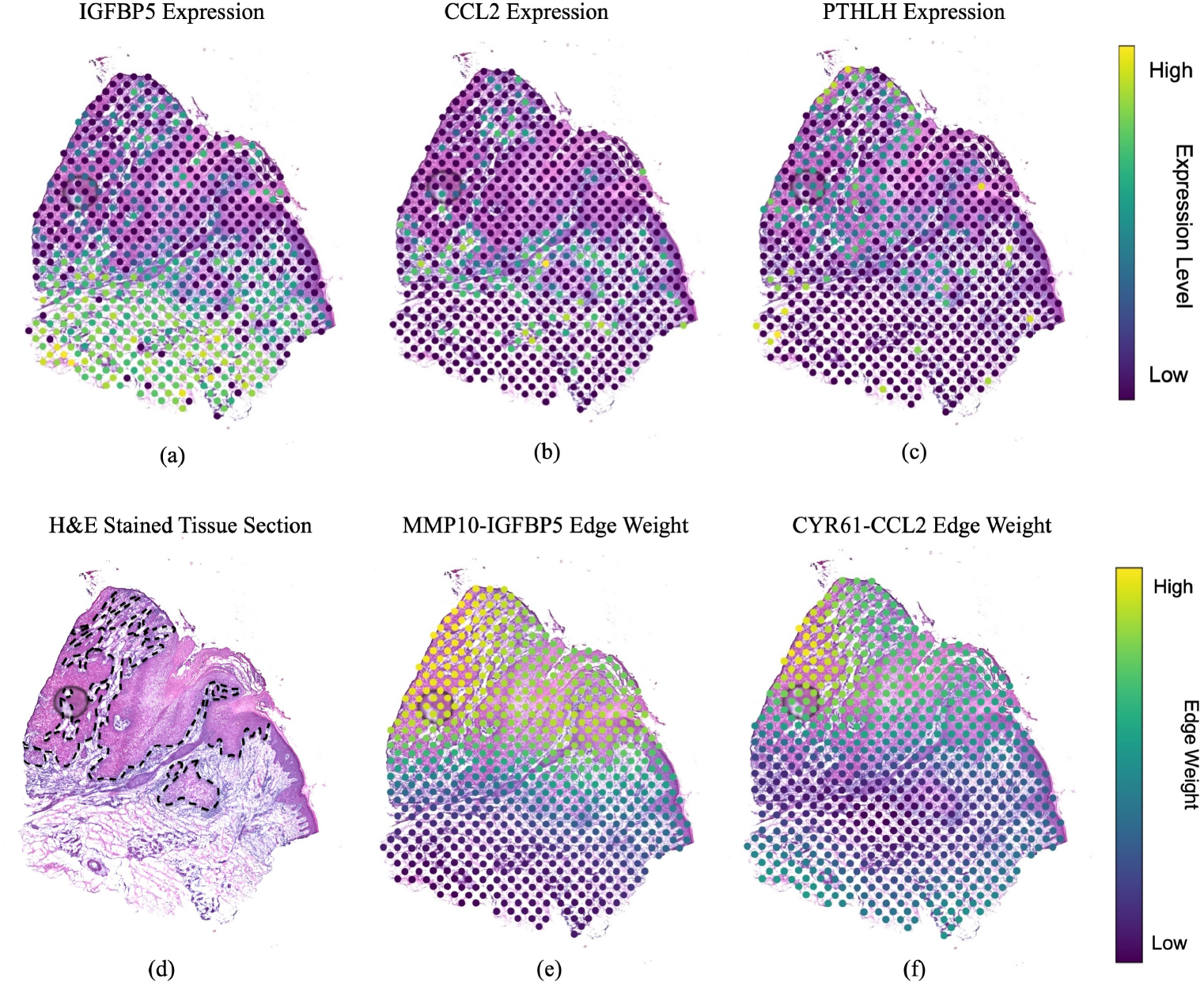
(a)-(c) Gene expression patterns of genes most frequently appeared in predicted high variable edges. (d) The H&E stained tissue section in this experiment with leading edge of tumor manually annotated (dotted lines) (e)-(f) Edge weight changing patterns of highly variable edges.

From the ranked highly variable edges, we found that our model is able to discover genes with spatially distinct functions that play key roles in leading to the discrepancy between tumor and normal regions. The most frequently appeared genes in the top 200 predicted highly variable edges are found to include IGFBP5, PTHLH, and CCL2, which have appearance frequencies of 42, 38, and 20 respectively. These genes all exhibit region-specific expression patterns in the tissue, as shown in Fig. 6 (a)-(c), and are supported by current studies indicating their distinct functional roles in normal skin versus tumor regions. IGFBP5, with higher expression in normal regions, has been shown to suppress tumor growth, while its down-regulation in tumors promotes cancer progression [69,42]. CCL2, a chemokine shown enriched at the tumor’s leading edge, may drive tumor spread and pre-metastatic niche formation by transforming non-neoplastic epithelial cells into invasive ones [28]. PTHLH is a proteinaceous hormone reported to contribute to the pathogenesis of oral squamous cell carcinoma, which exhibits a higher expression level in cSCC tumors than adjacent normal tissues. Studies indicate that PTHLH stimulates SCC cell growth via an autocrine/paracrine manner [9] and is upregulated in SCC to control epithelial-mesenchymal interactions [11]. The apparent spatial expression variations of these genes are controlled by their corresponding dynamic gene regulations, resulting in the frequent occurrence of these genes in the high variable edges of our predicted GRNs.

We further analyzed the spatial weight patterns of highly variable edges to examine how gene interaction strengths differ across regions, offering insights into abnormal gene regulation in tumor development. Matrix metalloproteinases (MMPs) are zinc-dependent endopeptidases that degrade extracellular matrix (ECM) components and play critical roles in tumor progression, including growth, invasion, metastasis, angiogenesis, and cell survival. They facilitate proteolytic cleavage of IGFBP5, thereby influencing the impact of IGFBP5 has in cancer [69]. We explored the spatial distribution of the edge weight between MMP10 and IGFBP5 across all spots. As visualized in Fig. 6 (e), the interaction strength aligns with gene functions, with shifts corresponding to the spatial organization of tumor and normal regions.

Another noteworthy edge is between CCL2 and CYR61. Fig. 6 (f) shows a similar pattern, where interaction weights are stronger in tumor areas and gradually weaken in normal skin. Studies have shown that CYR61 interacts with CCL2 to promote localized inflammation in several diseases [17,10]. In SCC, CYR61 acts as a positive growth regulator [37], while CCL2 recruits macrophages and other immune cells to tumors, promoting tumor growth and proliferation [70,30]. Through CCL2 upregulation in tumor regions, CYR61 enhances immune cell recruitment and is in turn activated by them, creating a positive feedback loop that intensifies inflammation within tumors. These findings illustrate our model’s ability to capture spatially specific, tumor-associated gene interactions across regions.

From predicted spot-specific GRNs in real spatial transcriptomics datasets, our model demonstrates its ability to capture genes with region-specific functions that are key to understanding tumor growth and immune response. Additionally, the edge weights in predicted GRNs reveal how gene regulatory relationships shift with cellular position. Such findings highlight the utility of our model in uncovering location-dependent gene interaction mechanisms. This high-resolution GRN construction has important implications for studying development, tumor biology, and microenvironmental impacts on cell behavior, potentially advancing targeted therapies and personalized medicine.

### 3.4 SVGRN helps identify and understand functional tissue niches through GRNs

We applied SVGRN to the Visium fallopian tube sample from the HuBMAP Program [26], using the predicted spot-specific GRNs to better understand the cell functional identities and their layout in the tissue. This approach directly highlight differential gene regulation relationship accross cells, providing clearer insight into cell activities compared with traditional reliance on only gene expression levels and marker genes for cell type identification. The predicted GRNs for each spot were clustered into five groups using the K-means method, as shown in Fig. 7 (b). The clustering results reveal distinct features of groups that correspond to functional roles within the tissue.

**Fig. 7.**
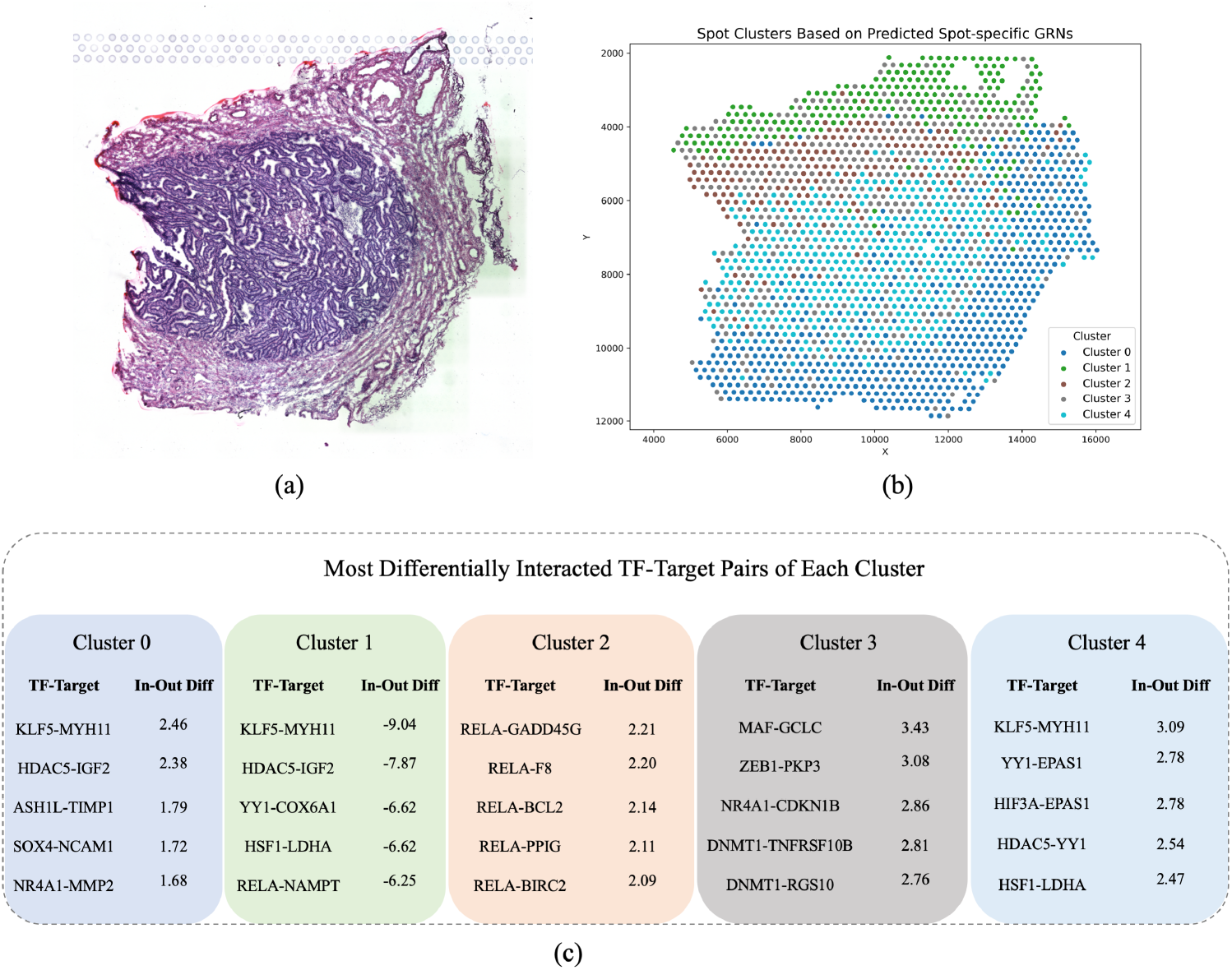
(a) The histology image of the fallopian tube sample. (b) Spots clustered by the predicted spot-specific GRNs. (c) The top differentially interacted TF-Target pairs in each cluster. *In-Out Diff* is the difference between the average interaction strength of spots in this cluster and out of this cluster, which are in 10^*−*3^ units.

To characterize the main differential cell activities, we obtained the possible TF-Target regulatory pairs from the TRRUST database [19], and calculated the average TF–Target interaction strength for each pair inside and outside each cluster. This allowed us to identify the most differentially interacted pairs. As shown in Fig. 7 (c), clusters 0, 2, 3, and 4 display the most differentially highly interacted pairs, while cluster 1 shows no stronger interactions than other clusters. Instead, it is defined by differentially weaker pairs. These top differential TF-Target regulation relationship reveals biological processes for each cluster and helps identify potential cell functions and tissue niches in the fallopian tube.

In cluster 0, the top interactions reflect cells near smooth muscle or fibroblasty regions at the epithelial boundary. KLF5 and MYH11 all play important roles in epithelial and smooth muscle cells [20,16], and NR4A1-MMP2 interactions occur in vascular smooth muscle cells [61]. IGF2 is involved in tissue regeneration in the fallopian tube after ovulation, promoting the transformation of fimbrial epithelial cells [22]. ASH1L–TIMP1 controls ECM remodeling, and NCAM1 contributes to cell–cell adhesion [31]. In contrast, cluster 1 shows distinct regulatory patterns. Many pairs that are strongly interacted in other clusters appear at lower strength, suggesting a possible identity of homeostatic epithelium which prioritizes ciliogenesis and beating over inflammatory, stress or ECM programs.

Cluster 2 highlights the regional inflammation repair processes. In the top interactions, the frequently appeared RELA regulates the inflammatory environment in the fallopian tube [66], GADD45 is associated with stress response, promoting the repair of DNA damage [24], and BCL2 supports cell survival in the inflammatory environment [39]. F8 plays a crucial role in the intrinsic pathway of blood coagulation [65] which may point cluster 2 toward a perivascular or endothelial neighborhood. Cluster 3 reflects calming signals, cleaning up, and rebuilding processes: GCLC protests against oxidative stress [38], NR4A1 and CDKN1B are involved in cell cycle control [21,56], and DNMT1-TNFRSF10B interaction is related to apoptosis thresholds tuning to remove damaged cells without excessive loss [75]. Cluster 4 represents a hypoxia-adapted perimuscular or epithelial interface. YY1-EPAS1 and HIF3A-EPAS1 indicate an active oxygen-sensing axis tuned to accommodate low-*O*_2_ niches [13], while HSF1-LDHA captures a heat-shock/proteostasis response coupled to a glycolytic shift typical of hypoxic remodeling.

Overall, these results highlight that SVGRN enables the identification of spatially distinct cell groups with unique GRN programs, which reflect their underlying functional identities and tissue roles. This GRN-based perspective provides a more mechanistic understanding of tissue organization, showing that functional niches in the fallopian tube can be uncovered not only by marker expression but also by the differential regulatory logic that drives cell activities.

### 3.5 Ablation Experiments

To assess the impact of our two key assumptions on cell-specific GRN learning, we conducted ablation experiments on two simulated datasets with noise levels of 0.1 and 0.5. In our final model, these assumptions are implemented by fixing each cell’s position to that of the target cell and by adding kernel weights to the loss function. In the ablations, we removed one or both of these components and compared the resulting average loss values, as shown in Table 1.

**Table 1.**
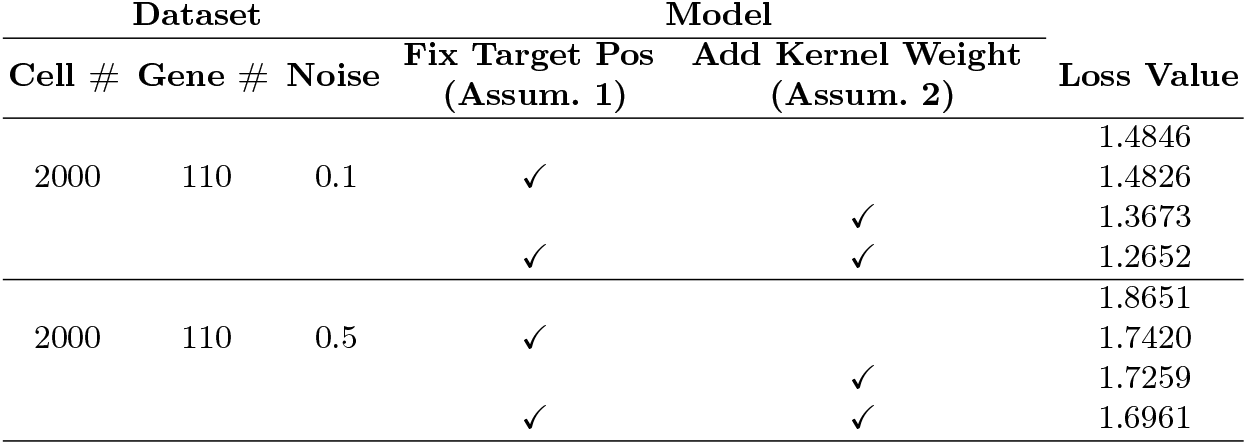
Ablation experiment results showing the changes in loss values when removing the assumption implement components in the model.

Without either assumption, the model cannot construct specific GRNs for each cell, resulting in identical GRNs across cells and the highest loss values in both datasets. As we incrementally reintroduce each assumption, we can observe significant loss reductions. This confirms that our assumptions not only allow for extending the whole-tissue GRN to single cells but also improve the reconstruction of gene expression based on spatial coordinates. By fixing coordinates alone, the model achieves a 0.13% and 6.6% loss reduction for noise levels 0.1 and 0.5, respectively. Adding kernel weights based on cell distances results in even greater reductions of 7.9% and 7.4% in the two cases.

## 4 Conclusion

In this paper, we proposed SVGRN, a deep learning based method for inferring cell-specific GRNs in spatial transcriptomics. It utilizes the gene expression data and spatial coordinates of cells to learn the underlying gene interaction in a self-regression way. The model is based on two assumptions about the relationship of a cell’s position to its GRN and the distribution of all GRNs in the space. These assumptions enable constructing the GRN for a single cell conditioned on its position and borrowing usable information from the cells nearby. From the experiments on simulated dataset, SVGRN shows advantages in more noisy and complex situations with more genes or cells considered. It also illustrates its ability in learning the changing gene regulation pattern from spatial transcriptomics dataset with different technologies and resolutions, such as spot-based and cell-based data. The future potential research could fall on more detailed understanding the causal relationship in gene regulations and involving biological interactions in other levels, such as the cell-cell interaction.

### Code and Data availability

The source code of SVGRN and configurations for CeSpGRN in experiments are available at https://github.com/lyrrrr/SVGRN. The seqFISH dataset of mouse embrys used in section 3.2 can be downloaded from https://content.cruk.cam.ac.uk/jmlab/SpatialMouseAtlas2020/. The cSCC spatial transcriptomics dataset can be downloaded from Gene Expression Omnibus (GEO) (https://www.ncbi.nlm.nih.gov/geo/) with accession number GSE144240. The sample *P2_ST_rep1* is used for our experiments in section 3.3. The fallopian tube data in section 3.4 comes from the HuBMAP Program (https://hubmapconsortium.org) and the sample used for our experiment can be found at https://portal.hubmapconsortium.org/browse/dataset/fa7fbcd8ae9219225f5df25e8c5e994e.

